# JAG1-NOTCH4 Mechanosensing Drives Atherosclerosis

**DOI:** 10.1101/2020.05.15.097931

**Authors:** Celine Souilhol, Xiuying Li, Lindsay Canham, Hannah Roddie, Daniela Pirri, Blanca Tardajos Ayllon, Emily V Chambers, Mark J Dunning, Mark Ariaans, Jin Li, Yun Fang, Maria Fragiadaki, Victoria Ridger, Jovana Serbanovic-Canic, Sarah de Val, Sheila E. Francis, Timothy JA Chico, Paul C Evans

**Affiliations:** Department of Infection, Immunity and Cardiovascular Disease, INSIGNEO Institute for In Silico Medicine, and the Bateson Centre, University of Sheffield, Sheffield, UK; Department of Pharmacy, Affiliated Hospital of Southwest Medical University, Luzhou, Sichuan 646000, P.R. China; Sheffield Bioinformatics Core, Sheffield Institute of Translational Neuroscience, University of Sheffield, Sheffield, UK; Department of Medicine, Biological Sciences Division, University of Chicago, Chicago, Illinois, USA; BHF Centre of Regenerative Medicine, Department of Physiology, Anatomy and Genetics, University of Oxford, Oxford, UK

## Abstract

Endothelial cell (EC) sensing of fluid shear stress regulates atherosclerosis, a disease of arteries that causes heart attack and stroke. Atherosclerosis preferentially develops at regions of arteries exposed to low oscillatory shear stress (LOSS), whereas high shear regions are protected. We show using inducible EC-specific genetic deletion in hyperlipidaemic mice that the Notch ligands JAG1 and DLL4 have opposing roles in atherosclerosis. While endothelial *Jag1* promoted atherosclerosis at sites of LOSS, endothelial *Dll4* was atheroprotective. Analysis of porcine and murine arteries and cultured human coronary artery EC exposed to experimental flow revealed that JAG1 and its receptor NOTCH4 are strongly upregulated by LOSS. Functional studies in cultured cells and in mice with EC-specific deletion of *Jag1* show that JAG1-NOTCH4 signalling drives vascular dysfunction by repressing endothelial repair. These data demonstrate a fundamental role for JAG1-NOTCH4 in sensing LOSS during disease, and suggest therapeutic targeting of this pathway to treat atherosclerosis.

## Introduction

Blood flow generates mechanical shear stress that has profound effects on the function of blood vessels by altering the physiology vascular endothelial cells (EC)^1^. Low oscillatory shear stress (LOSS) promotes the initiation and progression of atherosclerosis, a disease characterised by the accumulation of cells, lipids and other materials in the arterial wall that can lead to unstable angina, myocardial infarction or stroke^2,3^. Most regions of the arterial tree are exposed to physiologically high shear stress (HSS), which promotes EC quiescence and protection from disease. However, branches and bends of arteries are exposed to complex blood flow patterns, which LOSS that promotes EC dysfunction and the initiation of atherosclerosis. Intriguingly, LOSS drives both angiogenesis and atherosclerosis, suggesting a common mechanism for these divergent vascular processes^1^.

The Notch pathway was discovered in *Drosophila* as a master regulator of cell fate decisions and the spatial organisation of tissues^4^. Canonical Notch signalling involves an interaction between a Notch transmembrane receptor and a canonical Notch ligand on a contacting cell, which causes proteolytic cleavage of the Notch receptor by γ-secretase. This causes release of the Notch intracellular domain (ICD) which subsequently localises to the nucleus to regulate transcription^5^. Mammals possess four Notch receptors and five ligands. Classic studies revealed that the interaction between NOTCH1 and DLL4 on adjacent EC establishes cellular identity (i.e. ‘tip’ versus ‘stalk’ cells) during angiogenesis^6^, and NOTCH1 and DLL4 are critical for arterial specification^7–9^ and for neovascularisation in ischemic tissues^10,11^. Notch activation requires mechanical force induced by endocytosis of ligand^12^ and recent studies found that NOTCH1 sensing of shear stress is important in arterial differentiation^13^, vascular homeostasis^14^ and protection from atherosclerosis^15^. However, the potential ability of the Notch system to sense and respond to LOSS during atherosclerosis initiation has not been studied. Moreover, the function of Notch ligands in atherosclerosis has not been previously analysed.

Here we analysed the function of endothelial *Jag1* and *Dll4* in atherosclerosis using conditional gene deletion approaches and observed that these Notch ligand genes have opposing effects on atherosclerosis. While *Jag1* enhances atherosclerosis specifically at regions of LOSS that are susceptible to atherosclerosis, *Dll4* exerted an atheroprotective effect. Mechanistic studies revealed that NOTCH4 is the dominant Notch receptor at sites of LOSS, and that JAG1-NOTCH4 signalling enhances atherosclerosis by repressing EC proliferative reserve. These data fundamentally advance our knowledge of the role of Notch signalling in interpreting shear stress signals to control EC function and have important implications for therapeutic targeting of the Notch pathway in atherosclerosis.

## Results

### Endothelial *Jag1* and *Dll4* have divergent effects on atherosclerosis

To delineate the roles of endothelial *Jag1* and *Dll4* in atherosclerosis, we generated EC-specific inducible knockout mice by crossing floxed strains with a transgenic strain expressing *CDH5^CreERT2/+^.* Mice aged 6 weeks were treated with Tamoxifen for 2 weeks in order to delete *Jag1* or *Dll4* from EC or to generate experimental controls, and were then treated with adenoviral-PCSK9 and exposed to high fat diet for 6 weeks to generate hypercholesterolemia and atherosclerotic lesions (Fig. 1a and Fig. 2a). Tamoxifen treatment of *Jag1^fl/fl^ Cdh5^CreERT2/+^* mice *(Jag1^ECKO^*) induced deletion of *Jag1* which was validated by qRT-PCR (Fig. 1b; residual *Jag1* mRNA likely due to expression in smooth muscle cells) and en face staining of the aorta (Fig. 1c). Endothelial deletion of *Jag1* did not lead to clinical manifestations and did not affect cholesterol levels (Supplementary Fig. 1a). In hypercholesterolaemic mice, lesion area was significantly reduced in *Jag1^ECKO^* compared to controls in the whole aorta (Fig. 1d and Fig. 1e). A sub-analysis revealed that *Jag1^ECKO^* caused a reduction in lesion area in the aortic arch but not in the descending aorta (Fig. 1e). Lesion area in the aortic root was also unaltered in *Jag1^ECKO^* compared to controls (Fig. 1f and Fig. 1g). Therefore, *Jag1* promotes atherosclerosis specifically at the aortic arch which is a region exposed to LOSS.

**Figure 1.**
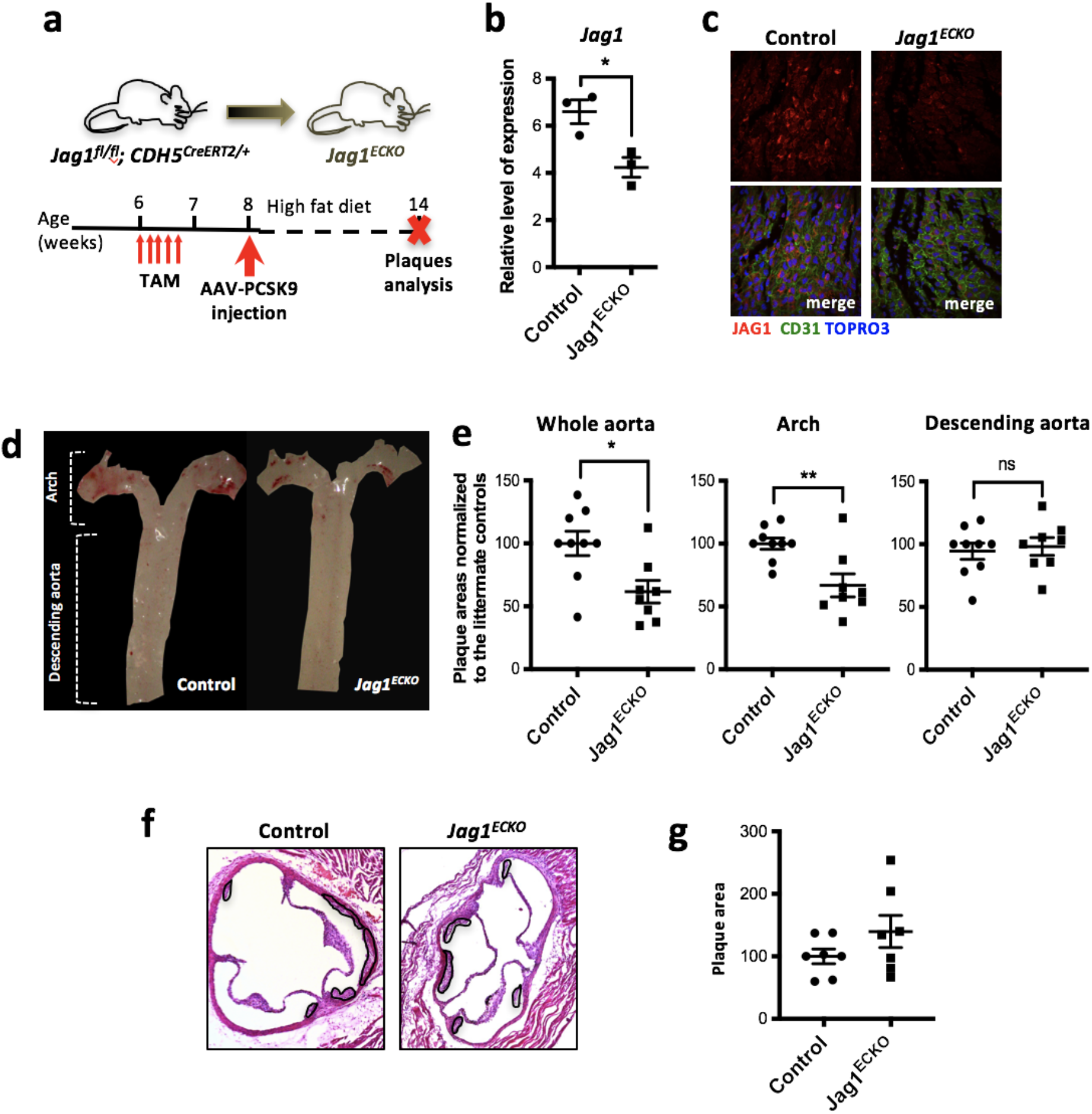
Loss of endothelial *Jag1* decreases plaque development at sites of low oscillatory shear stress. **(a)** Timeline of *Jag1* deletion in a model of hypercholesterolemia. *Jag1^fl/+^ CDH5^CreERT2/+^* (= *Jag1^ECKO^*) mice aged 6 weeks and littermates lacking Cre (*Jag1^fl/fl^*; = Controls) received five intraperitoneal injections of tamoxifen and one injection of PCSK9-AAV virus at specified time points. After 6 weeks fed with high fat diet, the mice were culled and plaque analysis performed by Oil Red O staining. **(b-c)** Validation of *Jag1* deletion. **(b)** Level of *Jag1* RNA determined by qRT-PCR in aortae isolated from *Jag1^fl/fl^ CDH5^CreERT2/+^* (= *Jag1^ECKO^* mice) (n=3) and littermate controls lacking Cre *(Jag1^fl/+^* or *Jag1^fl/fl^*; Controls) (n=3) two weeks post tamoxifen. **(c)** The expression of JAG1 protein (red) was visualized in the murine aorta by en face staining. EC were identified using anti-CD31 antibodies (green) and nuclei were costained using TOPRO-3 (blue). **(d)** Representative images of aortas stained with oil Red O and **(e)** normalized quantification of plaque burden in the whole aorta, arch, and descending aorta in *Jag1^ECKO^* mice (n=8) compared to littermate Controls (n=9). Representative images **(f)** and quantification of plaque burden **(g)** in the aortic roots of controls and *Jag1^ECKO^* mice. In all graphs, each data points represents one mouse and mean ± SEM are shown. Differences between means were analyzed using an unpaired *t*-test.

Deletion of *Dll4* was validated in aortae by qRT-PCR analysis in *Dll4^fl/fl^ Cdh5^CreERT2/+^* mice (*Dll4^ECKO^*; Fig. 2b). Deletion of both *Dll4* alleles caused lethality under high fat diet and we therefore analysed atherosclerosis using mice with a single allele deletion of *Dll4 (Dll4^ECHet^)*. En face staining using Oil-red-O revealed that lesion area in the aorta was significantly increased in *Dll4^ECHet^* compared to controls (Fig. 2c and Fig. 2d; Whole aorta). A sub-analysis of the aortic arch and descending aorta revealed that mean lesion area was enhanced in *Dll4^ECHet^* compared to controls at both sites but this difference reached statistical significance only in the aortic arch (Fig. 1d). Similarly, analysis of aortic root cross-sections revealed significantly enhanced lesion area in *Dll4^ECHet^* compared to controls (Fig. 1e and Fig. 1f). We also observed a slight increase of cholesterol and triglyceride levels in *Dll4^ECHet^* animals (Supplementary Fig. 1b). Thus, we conclude that endothelial *Dll4* exerts an atheroprotective function in the aorta and aortic root. In summary, endothelial *Dll4* and *Jag1* have divergent roles in atherosclerosis; *Dll4* is protective whereas *Jag1* initiates disease specifically at sites of LOSS.

**Figure 2.**
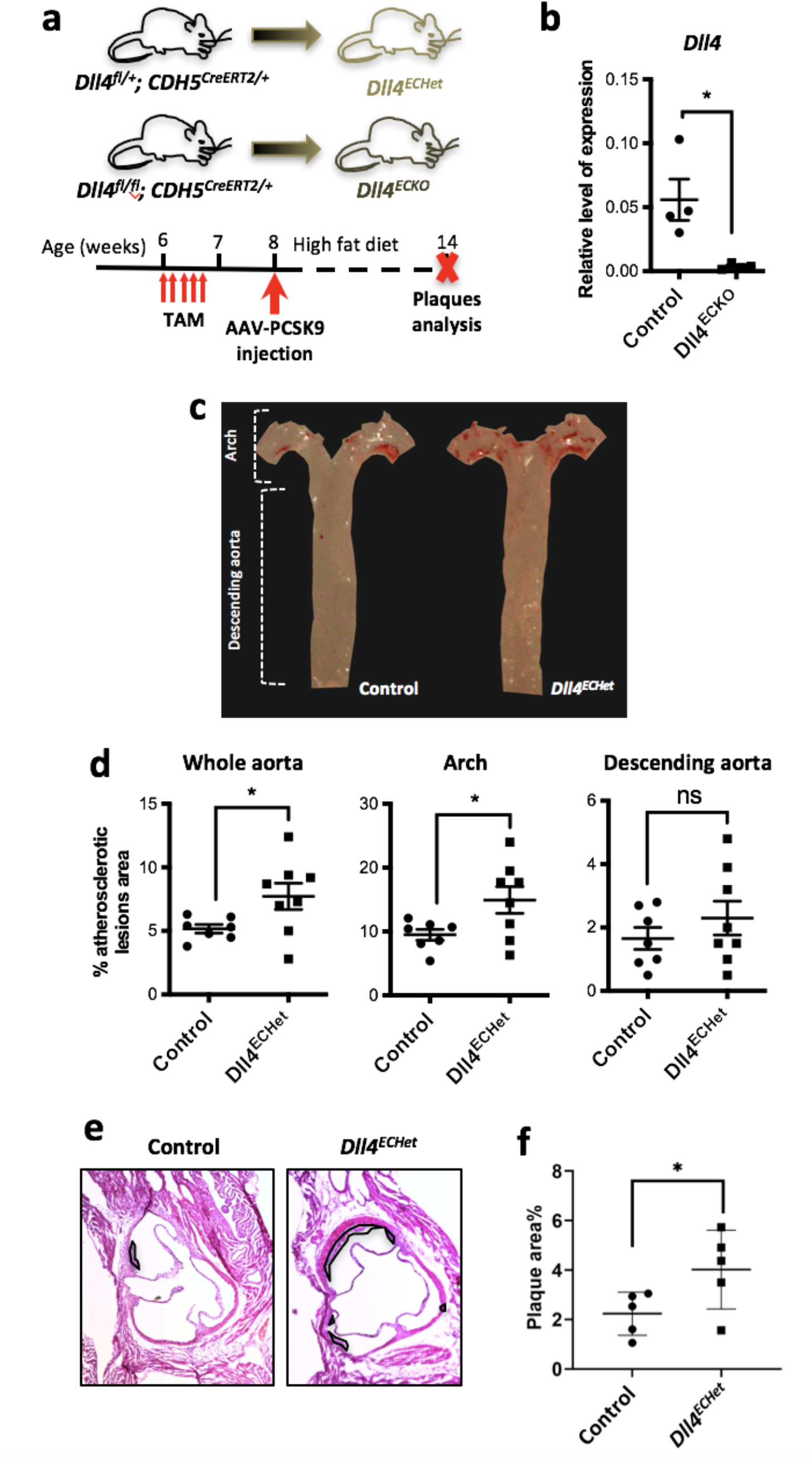
Loss of endothelial *Dll4* increases atherosclerotic plaque formation. **(a)** Timeline of *Dll4* deletion in a model of hypercholesterolemia. *Dll4^fl/+^ CDH5^CreERT2/+^* (= *Dll4^ECHet^*) and *Dll4^fl/fl^ CDH5^CreERT2/+^* (= *Dll4^ECKO^)* mice aged 6 weeks and littermates lacking Cre *(Dll4^fl/+^* or *Dll4^fl/fl^*; = Controls) received five intraperitoneal injections of tamoxifen and one injection of PCSK9-AAV virus at specified time points. After 6 weeks fed with high fat diet, the mice were culled and plaque analysis performed by Oil Red O staining. **(b)** Validation of *Dll4* deletion. *Dll4* transcript levels from aortae of adult control mice (n=4) and *Dll4^ECKO^* mice (n=4) two weeks post tamoxifen were measured by qRT-PCR. **(c)** Representative images of control and *Dll4^ECHet^* aortae stained with Oil Red O for quantification of atherosclerotic plaque area. **(d)** Quantification of plaque burden in the whole aorta, arch, and descending aorta was determined by calculating the percentage of aortic surface area covered by atherosclerotic lesions in each group (n = 7 for Control and n = 8 for *Dll4^ECHet^).* Representative images **(e)** and quantification of plaque burden **(f)** in the aortic roots of controls and *Dll4^ECHet^* mice. In all graphs, each data points represents one mouse and mean±SEM are shown. Differences between means were analysed using unpaired *t*-tests.

### JAG1 and NOTCH4 are enriched at atherosusceptible regions exposed to LOSS

Given the different effects of *Jag1* and *Dll4* on atherosclerosis, we hypothesised that Notch receptors and ligands may have a different spatial pattern of expression in arteries. This was tested by qRT-PCR analysis of transcripts of Notch receptors and their ligands isolated from regions of the porcine aorta exposed to LOSS (inner curvature) or HSS (outer curvature) using a shear stress map generated previously by our group^16^. The expression of *JAG1* and *NOTCH4* mRNA was significantly enhanced at sites of LOSS compared to HSS whereas the expression of *DLL4, NOTCH1, NOTCH2 and NOTCH3* was similar between these sites (Fig. 3a). Similarly, *en face* staining demonstrated that JAG1 and NOTCH4 proteins were expressed at higher levels at a LOSS region compared to a HSS region in the aorta of C57BL/6 mice (Fig. 3b and Fig. 3c). Moreover, endothelial expression of JAG1 was enhanced at atherosclerotic plaques compared to non-diseased arteries in hypercholesterolemic ApoE^-/-^ mice (Supplementary Fig. 2), suggesting a potential role for JAG1 in atheroprogression.

**Figure 3.**
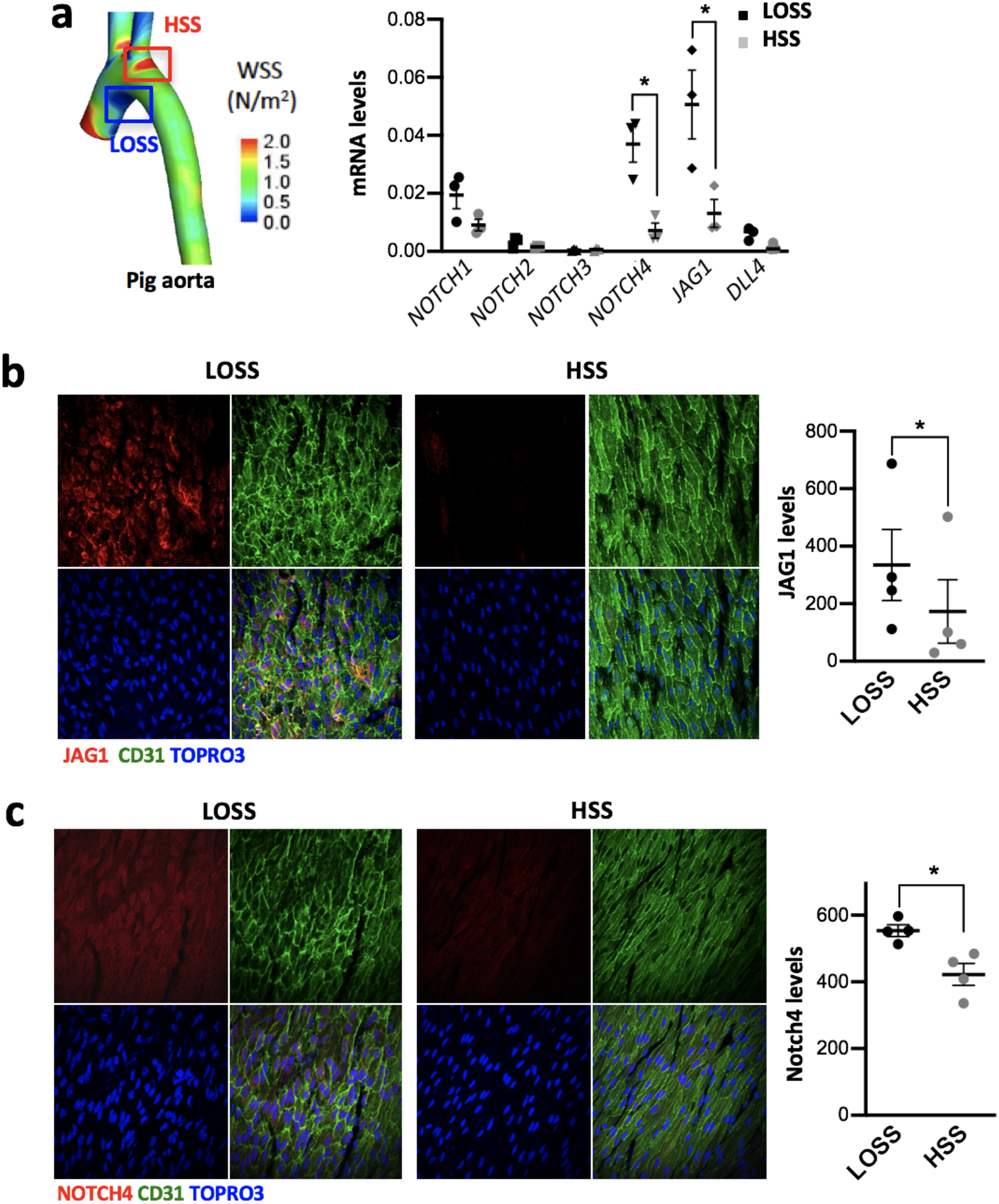
JAG1 and NOTCH4 are enriched at atheroprone sites. **(a)** MRI was used to measure geometries of the porcine dorsal aorta. Computational fluid dynamics simulation yielded a time-averaged WSS model of the aorta^16^. Based on this map, endothelial cells were specifically isolated from LOSS area versus HSS area in the porcine aortic arch. The expression of Notch actors was analysed in each population by qRT-PCR (n=3). **(b, c)** Aortic arches were isolated from C57BL/6 mice and en face immunostainings were performed using anti-JAG1 **(b)** or anti-NOTCH4 **(c)** antibodies (red). The endothelium was stained with anti-CD31 antibodies (green) and co-stained with TO-PRO-3 (DNA; blue). The graphs on the right represent the red mean fluorescence intensity (MFI) (n=4). Differences between means were analysed using paired *t*-tests.

### LOSS activates JAG1-NOTCH4 signalling

The correlation of JAG1 and NOTCH4 with LOSS does not reveal causality since atheroprone sites are exposed to alterations in mass transport, oxygen levels and inflammation as well as shear stress. We therefore used *in vitro* and *in vivo* models to assess directly whether Notch receptors and ligands are shear stress sensitive. A commercial syringe-pump, parallel plate apparatus was used to expose cultured human coronary artery EC (HCAEC) to either HSS or LOSS for 72 h. We validated the approach, by demonstrating that inflammatory *MCP1* transcripts are enhanced under LOSS whereas protective *KLF4* mRNAs are enhanced by HSS (Fig. 4a). Analysis of Notch pathway components by qRT-PCR revealed that *JAG1, DLL4, NOTCH4* and the Notch target gene *HES1* were significantly enhanced under LOSS compared to HSS, whereas *NOTCH1, NOTCH2 and NOTCH3* were unaltered (Fig. 4b). We next assessed the *activity* of Notch receptors using antibodies that bind specifically to the active, cleaved intracellular domains (ICD) of NOTCH4 (N4ICD) and NOTCH1 (N1ICD). It was observed by immunostaining (Fig. 4c) and immunoblotting (Fig. 4d) that N4ICD was significantly enhanced under LOSS compared to HSS, whereas N1ICD was not altered by shear stress (Fig. 4d). It should also be noted that detection of N1ICD required exposure of immunoblots for much longer periods compared to detection of N4ICD, suggesting that N4ICD is the dominant Notch receptor under LOSS (Fig. 4d). Immunoblotting also revealed that JAG1 was enhanced by exposure of HCAEC to LOSS, whereas DLL4 did not exhibit change at the protein level (Fig. 4d). Similarly, *in vivo* studies of JAG1 demonstrated that it is enhanced by LOSS in murine carotid arteries modified with a constructive cuff for 14 days (Fig. 4e).

At a mechanistic level, ATACseq analysis of HAEC exposed to flow revealed two regions in the *NOTCH4* gene where signals were enhanced under LOSS compared to HSS (Fig. 4f and Supplementary Fig. 3; Regions 1 and 2). Mining of published ChIPseq data from cultured EC^17,18^ revealed that these regions co-precipitate with ETS1 a master regulator of endothelial promoters and are associated with histone H3 lysine 4 acetylation (H3K4Ac) which marks genes for expression. Moreover, scanning of regions 1 and 2 using *Find Individual Motif Occurrences (FIMO)* identified putative consensus sequences for the shear stress sensitive transcription factors SMAD^19^, TWIST1^20^, Snail^21^ and NF-κB^22^ families (Supplementary Fig. 3). We conclude that *NOTCH4* is induced by LOSS via chromatin changes associated with shear-sensitive transcription factors. By contrast, ATACseq signals from the JAG1 gene were not altered by shear stress conditions (Supplementary Fig. 3) indicting that LOSS induction of JAG1 may not involve chromatin alteration. Collectively, these data reveal that JAG1 and NOTCH4 are the dominant components of the Notch pathway under conditions of LOSS.

### LOSS induces JAG1-dependent activation of NOTCH4

We investigated whether JAG1 is required for NOTCH4 activation by LOSS. The ability of ligands to elicit Notch signalling in response to LOSS was tested by applying anti-DLL4^23^ or anti-JAG1^24^ blocking antibodies and measuring N4ICD as an indicator of NOTCH4 activity. Of note both anti-JAG1 and anti-DLL4 blocking antibodies attenuate Notch activity level as revealed by a decreased expression of Notch target genes such as *HES1, HEY1* and *HEY2* (Supplementary Fig. 4). It was observed that blocking antibodies directed against JAG1 significantly reduced N4ICD levels whereas blocking of DLL4 activity had modest effects that did not reach significance (Fig. 5a). Thus, LOSS activation of NOTCH4 requires JAG1. Conversely, the induction of JAG1 by LOSS was abolished by treating HCAEC with the DAPT (an inhibitor of Notch receptor activity; Fig. 5b) indicating the Notch activity drives JAG1 expression. Thus, it was concluded that JAG1 and NOTCH4 interact at a functional level in endothelium exposed to LOSS.

**Figure 4.**
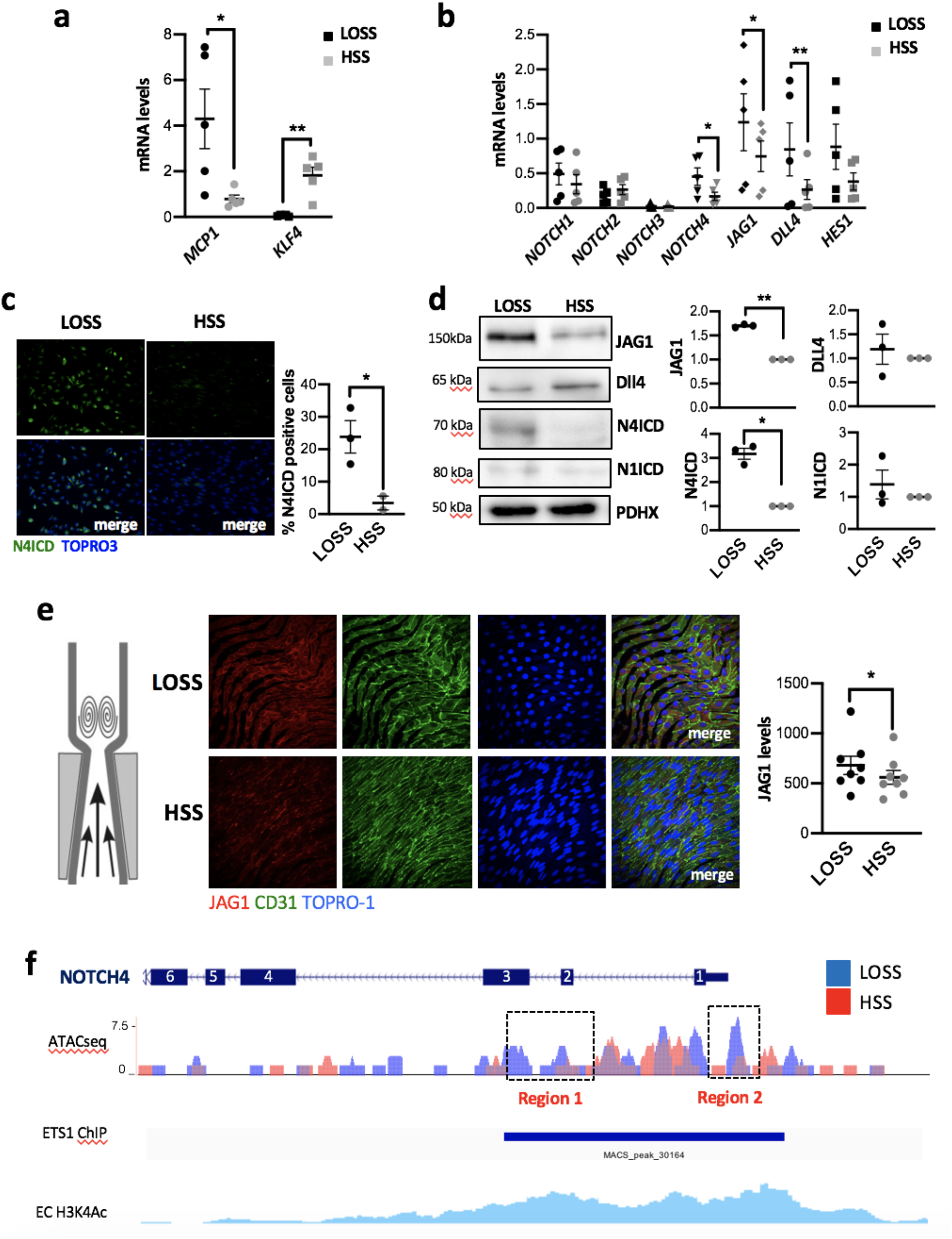
JAG1-NOTCH4 pathway is induced by LOSS *in vitro.* **(a, b)** Human coronary aortic endothelial cells (HCAEC) were seeded on **m** -slides and cultured under LOSS (4 dyn/cm^2^ oscillating at 1Hz) or HSS (13 dyn/cm^2^) for 72h using ibidi system. Expression level of known shear-sensitive genes (*MCP1* and *KLF4*) **(a)** or Notch actors **(b)** in HCAEC exposed to LOSS versus HSS for 72h was assessed by qRT-PCR. (n=5). **(cd)** Protein levels of Notch ligands and activated forms of NOTCH 1 and 4 receptors (N1ICD and N4ICD) in HCAEC exposed to LOSS and HSS for 72-96h were assessed by immunostaining (**c**: for N4ICD) (n=1) and western blot analysis (**d**: for DLL4, JAG1, N4ICD and N1ICD with normalization to the level of housekeeper gene PDHX) (n=3). Representative images are shown. **(e)** Flow-altering, constrictive cuffs were placed on the right carotid arteries of C57BL/6 mice. They generated anatomically distinct regions exposed to HSS and LOSS. Carotid arteries were harvested after 14 days, and en face staining was performed using anti-JAG1 antibodies (red), anti-CD31 antibodies (green), and the nuclear counterstain TO-PRO-3 (blue). Representative images and quantification of JAG1 expression are shown. (n=8 mice). Differences between means were analysed using paired *t*-tests. (f) HAEC were exposed to LOSS or HSS for 24 h prior to ATACseq and analysis of the *NOTCH4* gene. Regions showing enhanced accessibility under LOSS compared to HSS are identified (boxes; Region 1 and Region 2). Data were integrated with ETS1 and H3K4Ac ChIPseq analysis of human umbilical vein EC^17,18^.

**Figure 5.**
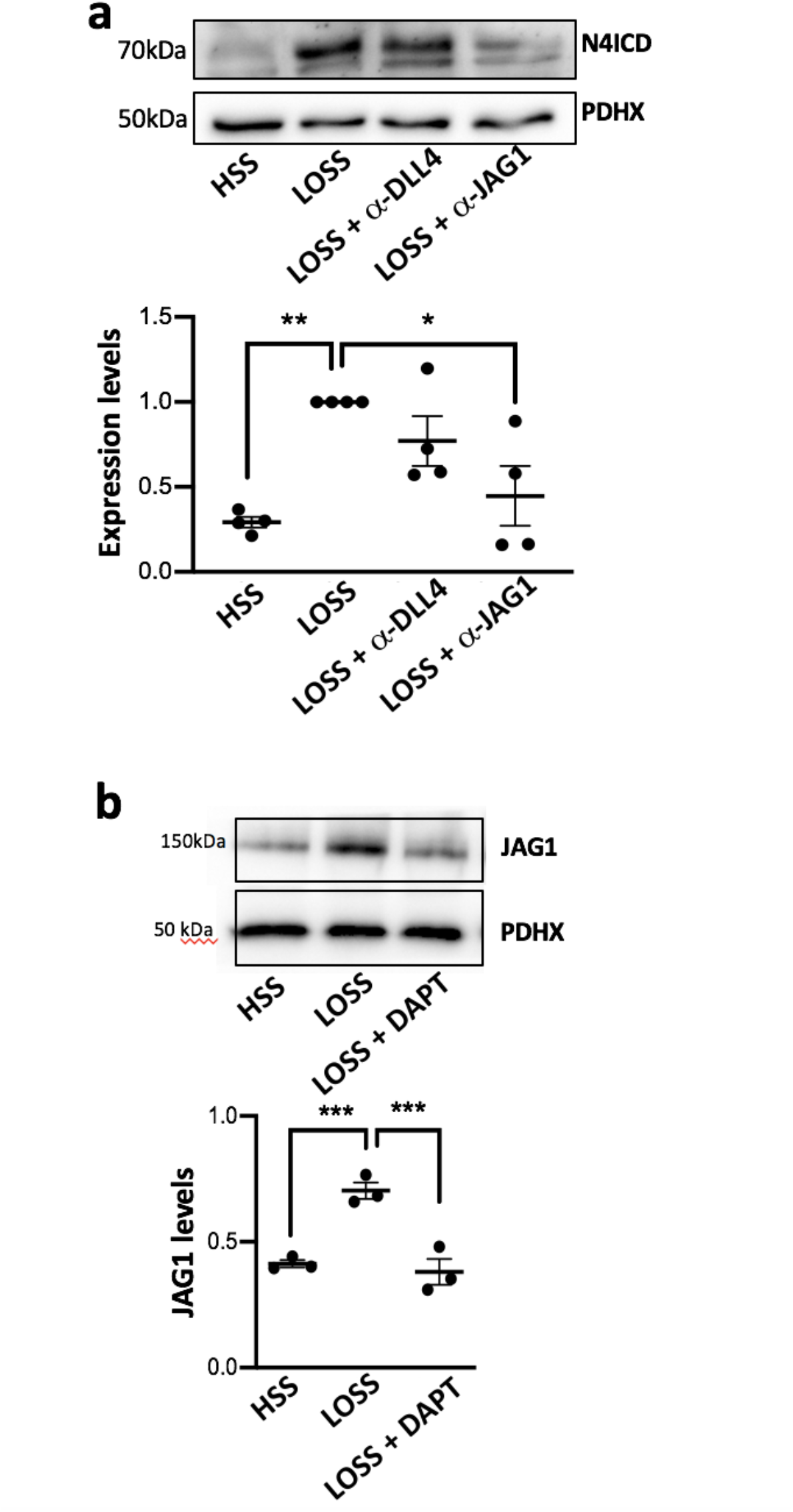
LOSS activation of NOTCH4 is Jag1-dependent. Human coronary aortic endothelial cells (HCAEC) were cultured under LOSS (4 dyn/cm^2^ oscillating at 1Hz) or HSS (13 dyn/cm^2^) for 72h using the Ibidi system either in the presence or absence of DAPT (γ-secretase inhibitor) or blocking antibodies against JAG1 or DLL4. Protein levels of the activated form of NOTCH4 (N4ICD) **(a)** or JAG1 **(a)** were assessed by western blot analysis and normalized to the level of housekeeper gene (n=3-4). Representative images of the western blots are shown for each gene. Differences between means were analysed using ANOVA test.

### JAG1 sensing of LOSS represses endothelial repair at atherosusceptible sites

The function of JAG1 was investigated by applying anti-JAG1 blocking antibodies to HCAEC exposed to LOSS for 48 h and quantifying changes in the transcriptome by RNAseq. The expression of 1239 genes was significantly altered by inhibition of JAG1, and functional annotation using DAVID found that the most highly enriched gene ontology terms for the genes negatively regulated by Jag1 are centred around cell division processes including Mitotic Nuclear Division, Cell Division, Positive Regulation of Mitotic Transition, cytokinesis (Fig. 6a and Table 1). The expression of JAG1-regulated genes was visualised using a volcano plot with the position of cell division regulators superimposed (Fig. 6b). This unbiased assessment of JAG1-regulated genes led to the hypothesis that JAG1-NOTCH4 signalling is a central negative regulator of EC proliferation. We confirmed this by demonstrating that HCAEC proliferation under LOSS is significantly enhanced by inhibition of Notch signalling (DAPT; Fig. 6c), by inhibition of JAG1 (blocking antibodies; Fig. 6d) or by *NOTCH4* knock-down (siRNA; Fig. 6e, Supplementary Figure 5). Notably, inhibition of JAG1 led to significantly increased EC proliferation in wounded monolayers exposed to LOSS (Fig. 6f), demonstrating that JAG1-NOTCH4 signalling reduces the capacity for endothelial repair. These observations were confirmed *in vivo* by en face staining of Ki67 which demonstrated that EC proliferation was significantly higher in *Jag1^ECKO^* compared to control mice (Fig. 6g). Collectively, these data indicate that JAG1 reduces the capacity of the endothelium for repair under LOSS by limiting EC proliferation.

**Figure 6.**
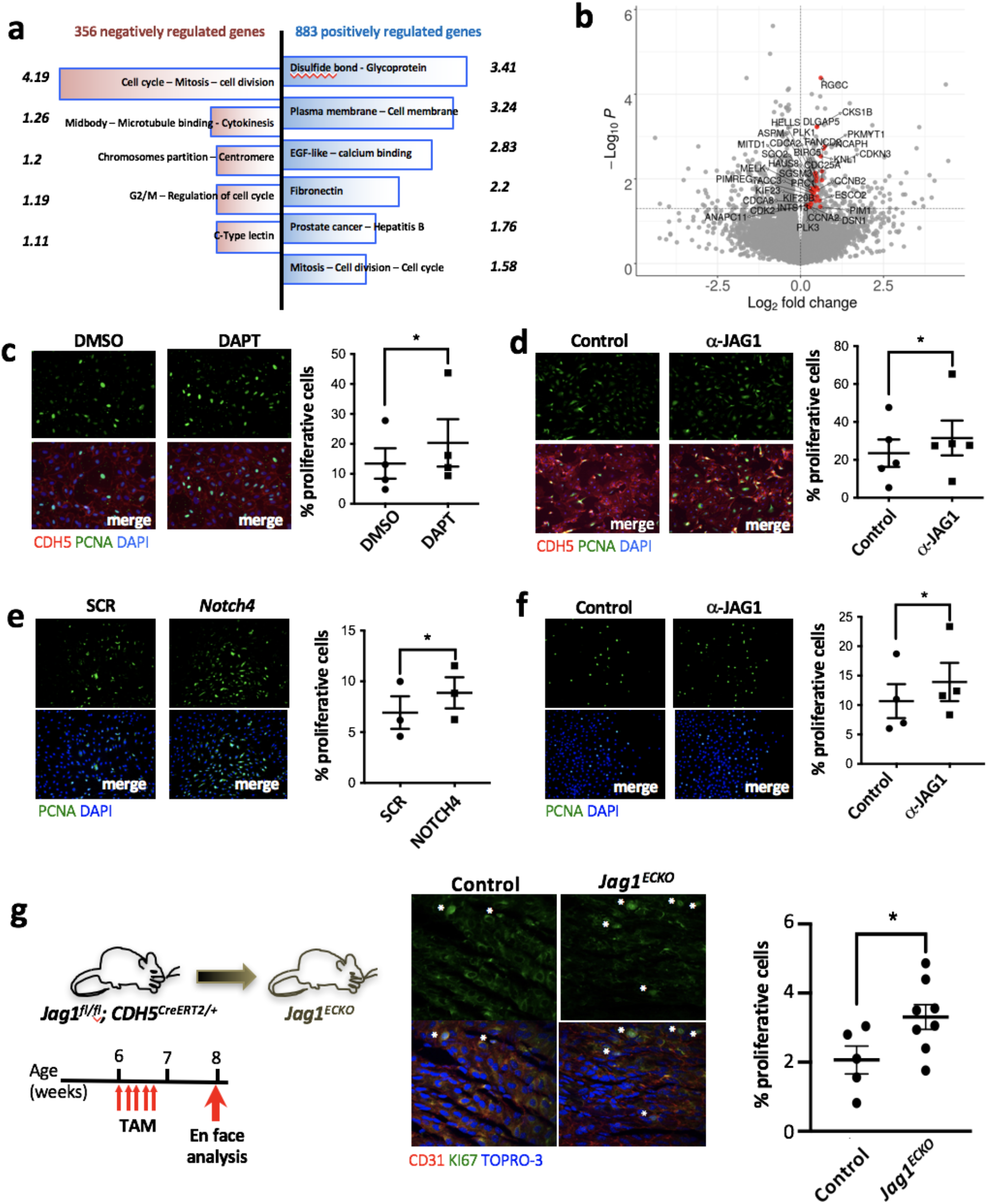
JAG1 activation by LOSS leads to inhibition of endothelial repair. **(a, b)** HCAEC isolated from 4 donors were exposed to LOSS and treated with **✓**-JAG1 blocking antibody for 48h before analysis for transcriptional changes by RNAseq. **(a)** Functional annotation clustering of the genes negatively (in red) and positively (in blue) regulated by JAG1 (p<0.05) using *DAVID bioinformatics resources.* Only the clusters with the highest enrichment score (indicated opposite each cluster) are shown. **(b)** Genes associated with cell cycle are upregulated following blockade with **✓**-JAG1. Volcano plot displaying differential expressed genes between JAG1 blockade and control samples. The y-axis corresponds to the mean expression value of -log10(pvalue), and the x-axis displays the log 2 fold change value. Significantly (p < 0.05) differentially expressed genes with a functional enrichment for the cell cycle are labeled and highlighted in red. **(c, d, e)** HCAEC were treated with either DAPT (γ-secretase inhibitor) **(c)** or **✓**-JAG1 blocking antibodies **(d)** or *NOTCH4* siRNA while exposed to LOSS for 72h using the Ibidi system. Proliferation was quantified by immunofluorescence staining using antibodies against PCNA (in green). EC were stained with CDH5 in red **(d, e)**. (n=3-5). **(f)** HCAEC were exposed to LOSS for 72h. To assess the role of JAG1 during endothelium repair, a scratch wound was made in the monolayer and the cells were treated with JAG1 blocking antibodies for 24h. Proliferation rate at the edge of the wound was then tested by using pCNA immunostaining (in green). **(c-f)** Differences between means were analysed using paired *t*-tests. **(g)** Endothelial cell proliferation at a LOSS region of the aorta (inner curvature of arch) was quantified in control (n=5) versus *Jag1^ECKO^* (n=8) mice 2 weeks post-tamoxifen (TAM) injection by en face immunostaining of the endothelium using antibodies against Ki67 (in green). EC were identified by using CD31 (in red) and nuclei were co-stained with TO-PRO-3 (in blue). Differences between means were analysed using unpaired T-test.

**Table 1.** JAG1 regulated genes in arterial endothelium.

|  | GO Categories | P Value |
| --- | --- | --- |
| **<u>356 negatively regulated genes</u>** | **Mitotic nuclear division**<br>**Cell division**<br>**Positive regulation of mitotic transition**<br>Angiogenesis | **(*p*=1.3E-4)**<br>**(*p*=2E-3)**<br>**(*p*=7.7E-3)**<br>(*p*=9.8E-3) |
| **<u>883 positively regulated genes</u>** | Angiogenesis<br>Positive regulation of intracellular signal transduction<br>**Regulation of endothelial cell proliferation**<br>Axon guidance<br>Signal transduction<br>Positive regulation of transcription<br>Regulation of chemokine biosynthesis<br>Lipopolysaccharide-mediated signaling pathway<br>Negative regulation of osteoblast differentiation<br>Regulation of excitatory postsynaptic potential<br>Cell adhesion<br>Positive regulation of ERK1/2 cascade | (*p*=6.6E-4)<br>(*p*=1.7E-3)<br>**(*p*=1.8E-3)**<br>(*p*=1.9E-3)<br>(*p*=2.0E-3)<br>(*p*=2.4E-3)<br>(*p*=3.3E-3)<br>(*p*=5.9E-3)<br>(*p*=6.3E-3)<br>(*p*=6.9E-3)<br>(*p*=7.3E-3)<br>(*p*=9.6E-3) |

The most significant functionally enriched gene ontology (GO) terms identified by DAVID are shown.

## Discussion

### DLL4 and JAG1 have divergent functions in atherosclerosis

Blood flow regulates multiple signalling pathways that control both angiogenesis and atherosclerosis^1^. The Notch pathway has a classical role in developmental angiogenesis^6^, arterial specification^7–9^ and neovascularisation^10,11^, and more recently has been characterised as a regulator of atherosclerosis^15,25–28^. Notch receptors and ligands are expressed in plaques^25,26^, and inhibition of *Dll4* or *Notch1* at a global level reduces atherosclerosis in murine models^25,27,28^. Moreover, tissue specific deletion of Notch receptors and ligands has revealed roles for this pathway in multiple cell types including macrophages^29^ and vascular smooth muscle cells^30^. Here we demonstrate that the Notch ligands DLL4 and JAG1 have opposing functions in atherosclerosis (Model: Fig. 7). Inducible deletion of *Dll4* from EC of hypercholesterolemic mice led to enhanced atherosclerosis. This function phenocopies *Notch1*^15^, and we therefore conclude that DLL4-NOTCH1 signalling is essential in limiting atherosclerosis. By contrast, we observed that JAG1-NOTCH4 signalling is an essential driver of atherosclerosis specifically at sites of LOSS. Therefore, we propose that the function of Notch signalling in atherosclerosis differs according to anatomical location; DLL4/NOTCH1 is atheroprotective at HSS regions, whereas JAG1/NOTCH4 induces disease at LOSS sites (Fig. 7). This model suggests that therapeutic targeting of JAG1/NOTCH4 signalling in EC may provide a novel treatment strategy to prevent or treat atherosclerosis.

**Figure 7.**
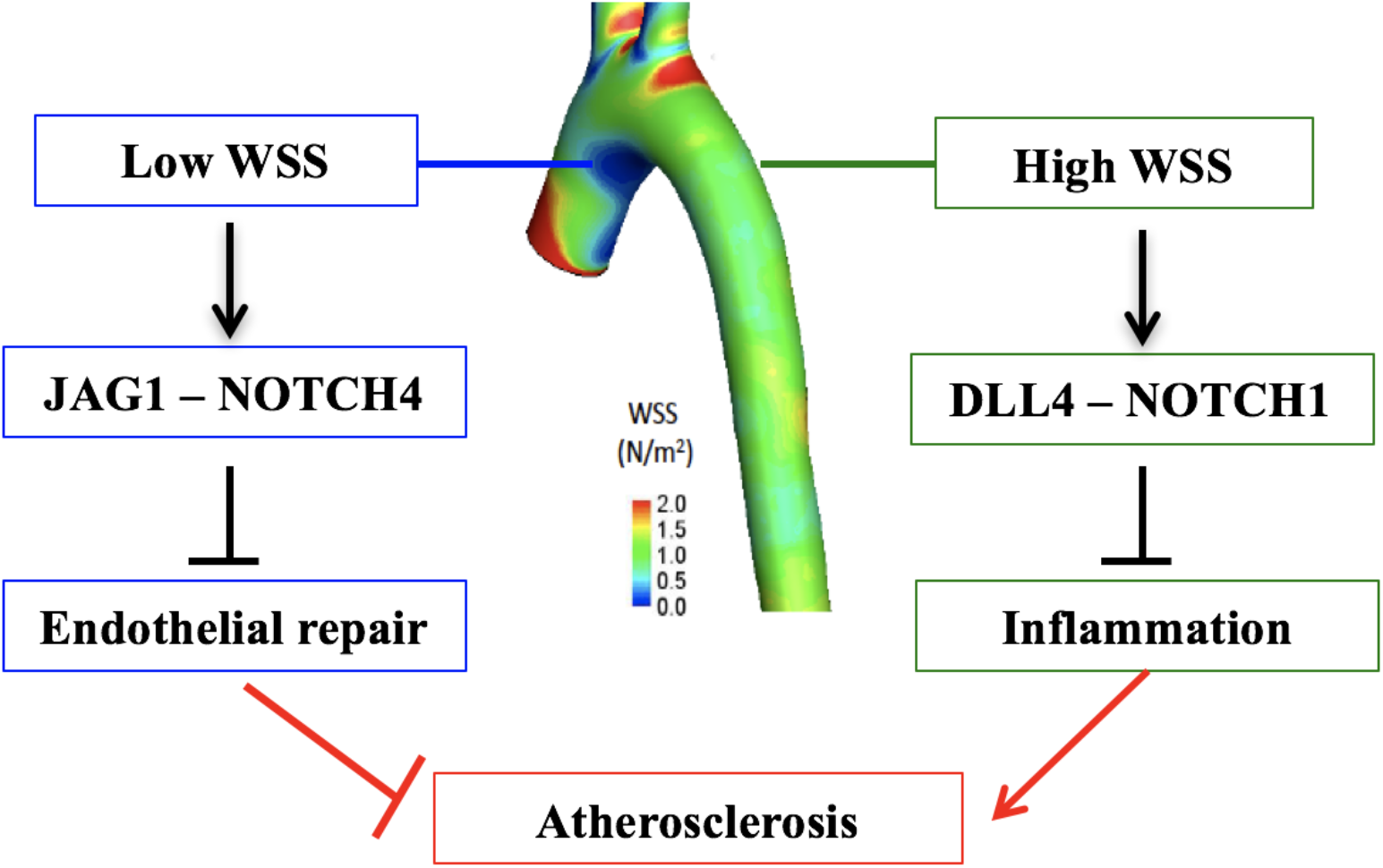
Proposed model for the divergent effects of JAG1-NOTCH4 signalling and DLL4-NOTCH1 signalling on endothelial function in atherosclerosis.

Using an unbiased RNAseq approach we show that the major function of JAG1/NOTCH4 in EC exposed to LOSS is to limit EC proliferation. This observation was validated experimentally by demonstrating that JAG1 and *NOTCH4* are negative regulators of proliferation in cultured EC exposed to LOSS, and that inducible deletion of *Jag1* rescues EC proliferation at sites of LOSS in the murine aorta. Several studies found that EC proliferation is enhanced at atherosusceptible LOSS regions of arteries^31,32^, however some have associated EC proliferation with disease processes^33^ whereas others suggest that it is atheroprotective^34^. We reconcile these different interpretations by proposing that EC proliferation can be both protective or disease-initiating depending on context. On one hand, the ability of EC to proliferate in response to injury at atheroprone sites is vital for vascular repair. Consistent with this, it has been demonstrated that irreversible cell cycle arrest (cellular senescence) of EC at atheroprone sites is an inducer of atherosclerosis^35^. On the other hand, uncontrolled proliferation of intact EC monolayers increases the permeability of arteries to lipoproteins and is therefore pro-atherogenic^33^. In summary, we propose that JAG1/NOTCH4 drives atherosclerosis by reducing proliferative reserve at atheroprone sites, thereby reducing the capacity for repair.

### JAG1/NOTCH4 mechanosensing

Sensing of mechanical force is fundamental to Notch signalling. In the canonical pathway, endocytosis of membrane bound ligand transmits force to the Notch receptor on a contacting cell leading to a structural change that promotes activation via γ-secretase cleavage^12^. Moreover, recent studies found that NOTCH1 is activated by physiological HSS. Polacheck et al demonstrated that physiological shear stress activates NOTCH1 to regulate the composition of adherens junctions and thereby control vascular permeability^14^. Consistent with this, Notch is also activated by physiological HSS during development to permit arterial differentiation^13,36,37^, cardiac morphogenesis^38,39^ and valve formation^40^ and is also activated by HSS in adult arteries to protect them from atherosclerosis^15^. The expression of JAG1 is also enhanced in HUVEC exposed to shear stress compared to static conditions^41^. While physiological HSS has been linked to Notch, we provide the first evidence that Notch signalling is also coupled to pathological disturbed blood flow patterns that drive vascular dysfunction and atherosclerosis. Our conclusion is based on the following observations: (1) JAG1 and NOTCH4 were exclusively enriched at LOSS regions of the aorta, (2) LOSS induces the expression of *NOTCH4* and *JAG1* and activates JAG1-NOTCH4 signalling, (3) JAG1-NOTCH4 signalling leads to defective endothelial repair and (4) inducible deletion of *Jag1* from murine EC reduced atherosclerosis specifically at LOSS regions of arteries. It remains to be determined whether NOTCH4 signalling responds to LOSS directly, i.e. mechanical force transduction through the Notch receptor and/or ligand, or whether the response is indirect involving other mechanoreceptors and further work is required to discriminate between these possibilities.

In summary, JAG1-NOTCH4 sensing of LOSS is a key driver of atherosclerosis by repressing transcriptional programmes that maintain EC proliferation and repair. These data demonstrate a fundamental role for JAG1-NOTCH4 in sensing disease-priming LOSS, and suggest that therapeutic targeting of endothelial JAG1-NOTCH4 could be a novel treatment strategy for atherosclerosis.

## Methods

### Mice

Mice with inducible deletion of *Jag1* or *Dll4* in EC were generated by crossing *Jag1^fl/fl^* ^42^ mice or *Dll4^fl/fl^* mice^43^ respectively with *CDH5^Cre-ERT2^* mice^*44*^. All mice were on a C57BL/6 background. PCR primers used for genotyping are described in Supplementary Table 1. To activate Cre, tamoxifen (Sigma) in corn oil was administered intraperitoneally (IP) for 5 consecutive days (2 mg/mouse/d). Two weeks after the first injection of tamoxifen, hypercholesterolemia was induced by intraperitoneal (I.P.) injection of adeno-associated virus containing a gain-of-function mutated version of proprotein convertase subtilisin/kexin type 9 (rAAV8-D377Y-mPCSK9) gene (Vector Core, North Carolina) followed by a high fat diet (SDS UK, 829100) for 6 weeks as previously described^45^. Constrictive cuffs were applied to the right carotid artery of anesthetized C57BL/6 mice as described previously^46^. C57BL/6 and transgenic mice were housed under specific-pathogen free conditions. Animal care and experimental procedures were carried out under licenses issued by the UK Home Office and local ethical committee approval was obtained. Mice between 1.5 and 3 months of age were used for experimentation.

### En face staining of murine endothelium

The expression levels of specific proteins were assessed in EC at regions of the inner curvature (LOSS site) and outer curvature (HSS site) of murine aortae or in the carotid by *en face* staining. Animals were killed by I.P. injection of pentobarbital and aortae were perfused *in situ* with PBS and then perfusion-fixed with 4% Paraformaldehyde (PFA) prior to harvesting. Fixed aortae were tested by immunostaining using specific primary antibodies (Supplementary Table 2). EC were identified by co-staining using anti-CD31 antibodies. Nuclei were identified using To-Pro-3. Stained vessels were mounted prior to visualization of endothelial surfaces *en face* using confocal microscopy (Olympus SZ1000 confocal inverted microscope). The expression of particular proteins at each site was assessed by quantification of the mean fluorescence intensities with standard error of the mean.

### EC culture and exposure to WSS

Human coronary artery EC (HCAEC) were purchased from PromoCell and cultured according to the manufacturer’s recommendations. HCAEC at passage 3-5 were seeded onto 0.4 mm Ibidi gelatin-coated μ-Slides (Luer ibiTreat, ibidi^TM^) and used when fully confluent. Flowing medium was then applied using the Ibidi pump system to generate low oscillatory (+/- 4 dyn/cm^2^, 0.5 Hz) or high (13 dyn/cm^2^) WSS. The slides and pump apparatus were placed in a cell culture incubator at 37°C. Inhibition of Notch activity was performed by addition of DAPT (50μM; Calbiochem) or blocking antibodies for Notch ligands (10μg/ml; anti-JAG1 or anti-DLL4) (Supplementary Table 2). Humanized phage antibody YW152F targeting DLL4^6^ was provided by Genentech.

### Immunofluorescent staining of cultured EC

HCAEC were fixed with PFA (4%) and permeabilised with Triton X-100 (0.1%). Following blocking with goat serum for 30 min, monolayers were incubated for 16 h with primary antibodies against PCNA (proliferation marker) and CDH5 (endothelial marker) (Supplementary Table 2) and AlexaFluor488- or Alexafluor568-conjugated secondary antibodies. Nuclei were identified using DAPI (Sigma). Images were taken with a widefield fluorescence microscope (LeicaDMI4000B) and analysed using Image J software (1.49p) to calculate the frequency of positive cells. Isotype controls or omission of the primary antibody was used to control for non-specific staining.

### *In vitro* repair assay

HCAEC were cultured until confluent in 6 well plates and exposed to LOSS using an orbital shaking platform housed inside a cell culture incubator and rotating at 210 rpm. This system generated low shear stress (approximately 5 dyn/cm^2^) with rapid variations in direction at the centre. After three days of culture under these conditions, wounds were created on confluent HCAEC monolayers using a pipette tip. Proliferation at the wound edge was assessed 24h later by immunofluorescence.

### Gene silencing

HCAEC cultures were transfected with siRNA sequences that are known to silence *NOTCH4* (L-011883-00, Dharmacon) using the Lipofectamine^®^ RNAiMAX transfection system (13778-150, Invitrogen) following the manufacturer’s instructions. Non-targeting scrambled sequences were used as a control (D-001810-01-50 ON-TARGETplus Non-targeting siRNA#1, Dharmacon).

### Isolation of EC from porcine aortae

Pig aortas from 4-6 months old animals were obtained immediately after slaughter from a local abattoir. They were cut longitudinally along the outer curvature to expose the lumen. EC exposed to high (= outer curvature) or low (= inner curvature) wall shear stress were harvested using collagenase (1 mg/ml for 10 minutes at room temperature) prior to gentle scraping.

### Real time PCR and RNAseq

RNA from mouse aortas was extracted after mechanical homogenization of the aorta in lysis buffer (Qiagen) using Triple-pure zirconium beads and a microtube homogenizer (Benchmark Scientific) at 4°C. RNA was extracted using the RNeasy Mini Kit (74104, Qiagen) and reverse transcribed into cDNA using the iScript cDNA synthesis kit (1708891, Bio-Rad). QRT-PCR was used to assess the levels of transcripts with gene-specific primers (Supplementary Table 3). Reactions were prepared using SsoAdvanced universal SYBR^®^Green supermix (172-5271, Bio-rad) and following the manufacturer’s instructions, and were performed in triplicate. Expression values were normalized against the house-keeping gene (mouse *Tbp,* human *HPRT* or porcine *B2M).* Data were pooled from at least three independent donors and mean values were calculated with SEM. For RNAseq, the purity and integrity of total RNA samples isolated from HCAEC was assessed using a Bioanalyser (Agilent) and high-quality samples from 4 HCAEC donors were used to prepare RNA-seq libraries that were sequenced on an Illumina HiSeq platform yielding 20 million reads per sample. Library preparation and cDNA sequencing were performed by Novogene. Fastq samples were processed using the RNA-seq pipeline implemented in the bcbio-nextgen project https://bcbionextgen.readthedocs.io/en/latest/index.html. After quality control checking using fastQC, RNA-seq reads were aligned to the human reference genome (assembly GRCh37/hg19) using STAR^47^ with the default parameters. FeatureCounts^48^ was used to create a matrix of mapped reads per Ensembl annotated gene. Differential gene expression was performed using the DESeq2 R package^49^. Functional enrichments for protein coding genes with p value < 0.05 and log2 fold change > 0 were calculated using DAVID^50^ using the total genes present as a background set.

### Western blotting

Total cell lysates were isolated using lysis buffer (containing 2% SDS, 10% Glycerol and 5% β-mercaptoethanol). Primary antibodies used and concentrations are described in Supplementary Table 2. HRP-conjugated secondary antibodies (Dako) and chemiluminescent detection was carried out using ECL Prime^®^ (GE Healthcare). Membranes were imaged using the Gel Doc XR+ system (Biorad).

### Lipid measurement

Blood samples were collected by terminal cardiac puncture and plasma was separated by centrifugation for further analysis using a COBAS analyser (total plasma cholesterol, non-HDL cholesterol and triglycerides).

### Atherosclerosis plaque analysis

Mice were euthanized and perfused-fixed with PBS followed by 4% PFA. The aorta was dissected, gently cleaned of adventitial tissue and stained with Oil Red O (Sigma). The surface lesion area was analysed using NIS elements analysis software (Nikon, NY). For analysis of aortic root sections, the upper portion of the hearts were dissected horizontally at the level of the atria and placed in 30% sucrose for 24 h before embedding in Optical Cutting Temperature (OCT). Serial 7 μm sections were processed for staining with Mayer’s hematoxylin. NIS-elements analysis software (Nikon, NY) was used to calculate the total lesion area.

### ATAQseq

ATAC-seq was performed on HAECs subjected to 24 h unidirectional flow or disturbed flow as previously described^51^.

## Statistical analysis

Data are presented as means values ± SEM. Statistical analysis were performed with GraphPad Prism software. The degree of significance is as following: *p <0.05; **p<0.01; ***p < 0.001. The test performed is indicated in the figure legend.

## Acknowledgements

This study was funded by the British Heart Foundation and also supported by the NIHR Sheffield Biomedical Research Centre / NIHR Sheffield Clinical Research Facility. We thank Professor Freddy Radtke for providing Dll4 and Jag1 floxed strains of mice.

## Author contributions

X.L., M.J.D., Y.F., M.F., V.R., J.S-C., S.D.V., S.E.F., T.J.A.C. contributed to the design of experiments, acquisition and analysis of data and preparation of the paper. L.C., H.R., D.P., B.T.A., E.V.C., M.A., J.L. contributed to the acquisition and analysis of data. C.S. contributed to the conception and design of experiments, acquisition and analysis of data and the preparation of the paper. P.C.E. conceived the study, designed the experiments, contributed to the acquisition of data, analysed the data and wrote the paper. All authors reviewed the paper and provided intellectual content

