## Supplementary Figures and Tables for "JAG1-NOTCH4 Mechanosensing Drives Atherosclerosis"

**Supplementary Figure 1**

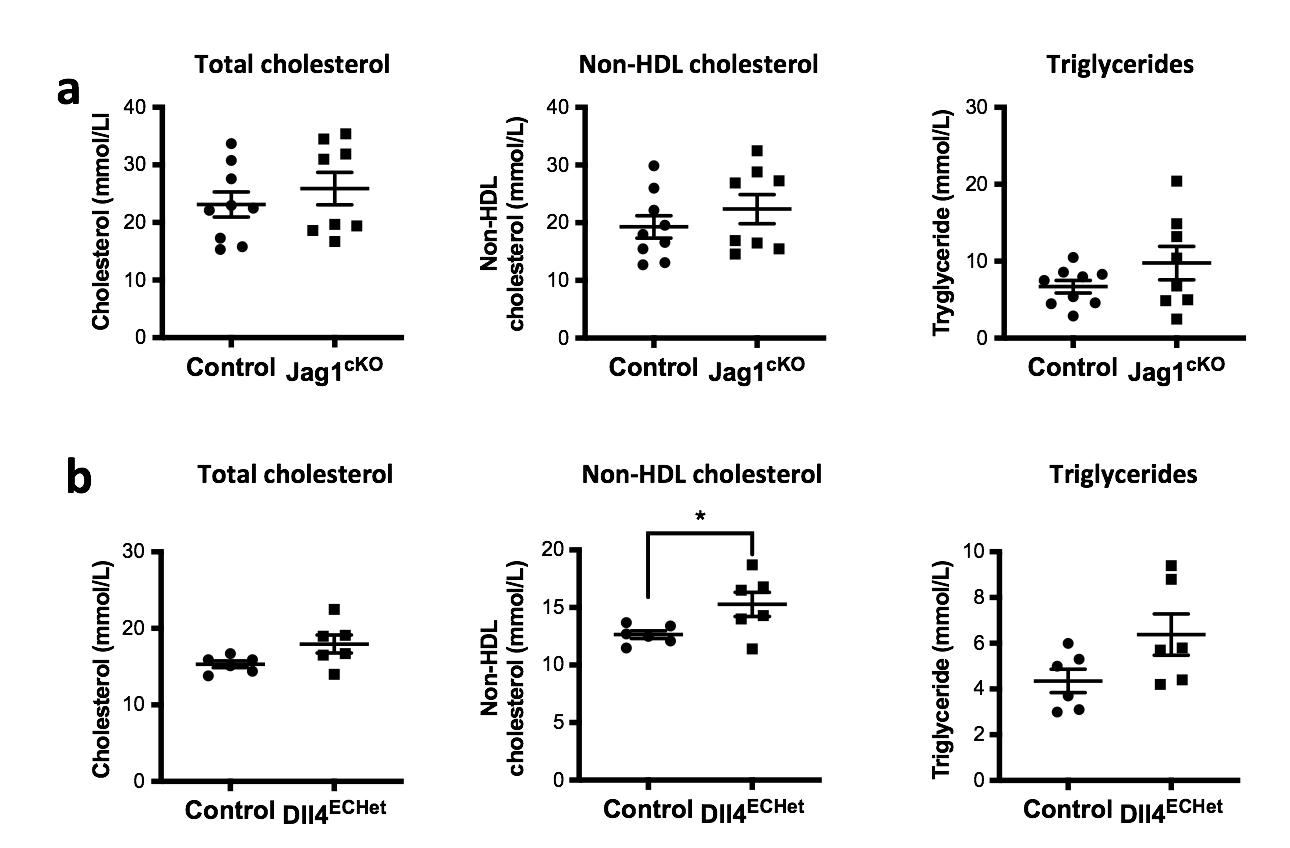

**Lipid profiles in mice with endothelial deletion of *Jag1* or *Dll4*.**

*Jag1^ECKO^* mice **(a)** or *Dll4^ECHet^* mice **(b)** aged 6 weeks and littermate controls received five intraperitoneal injections of tamoxifen and one injection of PCSK9-AAV virus at specified time points. After 6 weeks fed with high fat diet, the mice were culled and total cholesterol, non-HDL cholesterol and triglyceride levels were measured. Differences between means were analyzed using an unpaired t-test.

**Supplementary Figure 2**

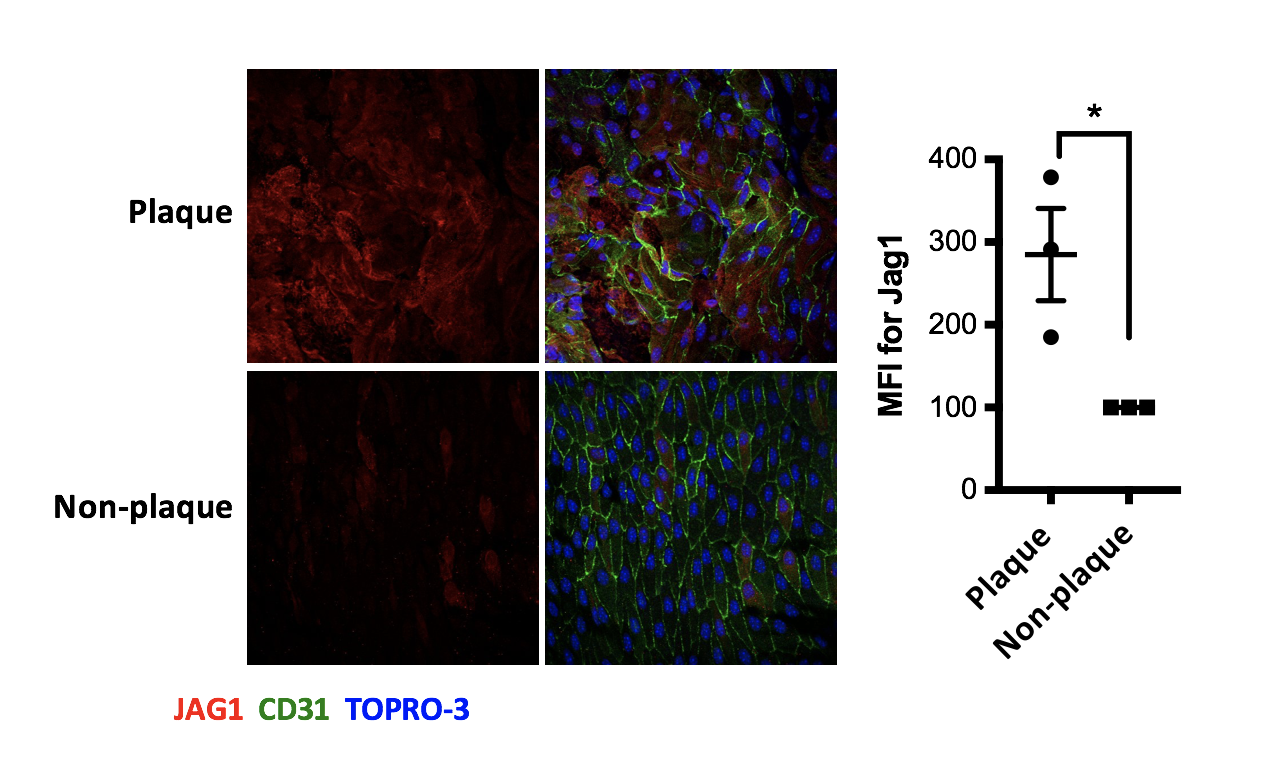

**JAG1 is expressed in endothelium overlying atherosclerotic plaques.**

*APOE^-/-^* mice were exposed to a high fat diet for 6 weeks prior to en face staining of the aorta using anti-JAG1 antibodies (red). The endothelium was stained with anti-CD31 antibodies (green) and co-stained with TO-PRO-3 (DNA; blue). Expression of JAG1 in the endothelium above atherosclerotic plaque versus non-plaque regions was quantified. Differences between means were analysed using paired t-tests.

**Supplementary Figure 3**

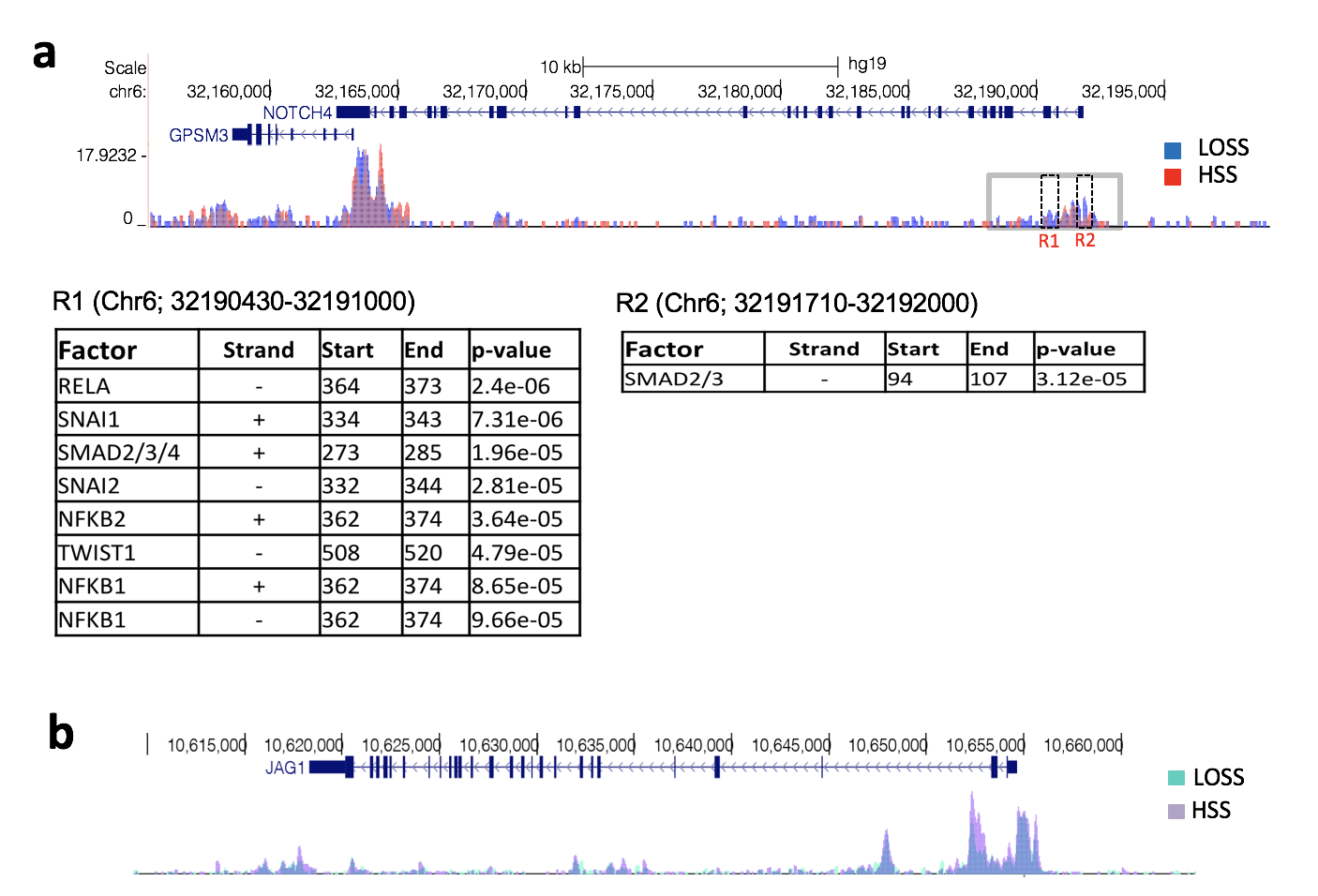

**ATACseq analysis of NOTCH4 and JAG1 in endothelial cells exposed to shear stress.**

ATAQseq experiments were conducted in HAEC exposed to 24h of LOSS or HSS. The results at *NOTCH4* **(a)** and *JAG1* **(b)** loci are depicted. **(a)** Sequences showing different accessibility under LOSS and HSS conditions (black boxes; region 1 (R1) and region 2 (R2)) were analysed using Find Individual Motif Occurrence (FIMO)^1^ in the MEME dataset to find transcription factors binding sites. The region boxed in grey is shown in detail in Figure 4f.

**Supplementary Figure 4**

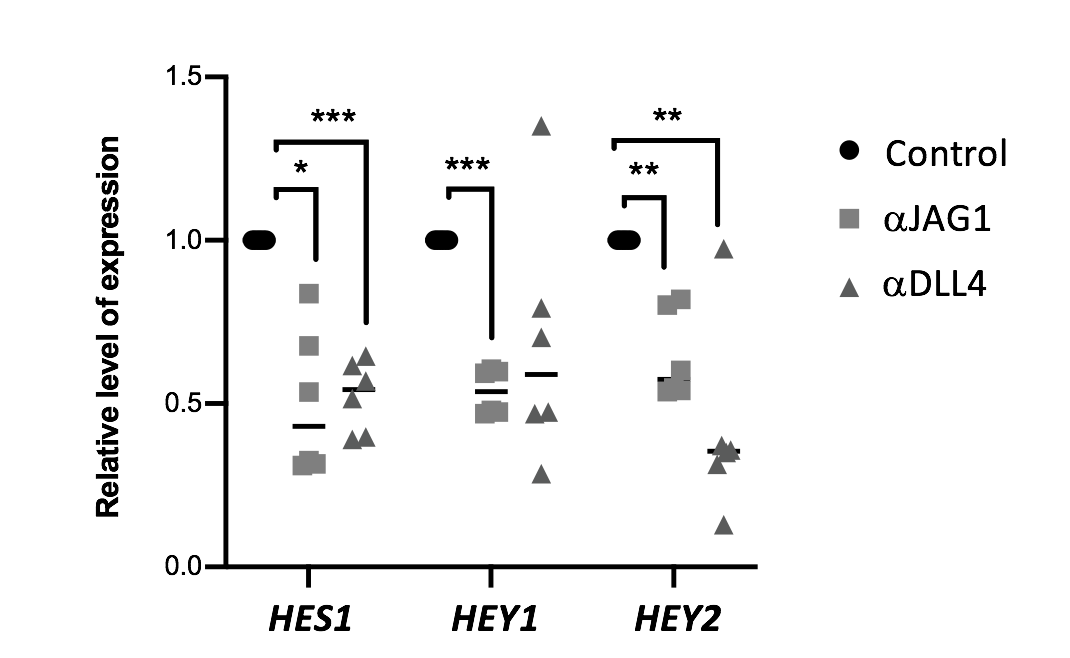

**Effect of DLL4 and JAG1 blocking antibodies on Notch target genes.**

HCAEC were cultured under LOSS (4 dyn/cm^2^ oscillating at 1Hz) or HSS (13 dyn/cm^2^) for 48h either in the presence or absence of blocking antibodies against JAG1 or DLL4. Expression levels of Notch target genes (HES1, HEY1, HEY2) were quantified by qRT-PCR. (n=6). Differences between means were analysed using one-way ANOVA.

**Supplementary Figure 5**

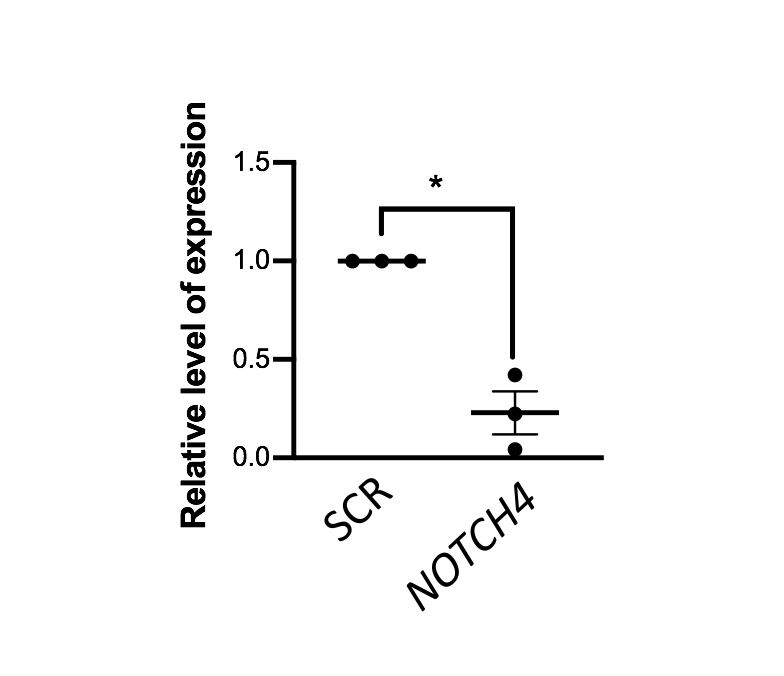

**Validation of *NOTCH4* siRNA.**

HCAEC were treated with *NOTCH4* or scrambled (SCR) control siRNA while exposed to LOSS for 48h using the Ibidi system. Expression levels of *NOTCH4* were quantified by qRT-PCR. (n=6). Differences between means were analysed using a paired t-test.

**Supplementary Table 1**

**PCR primers for mouse genotyping**

| **Gene** | **Forward** | **Reverse** | **Size** |
| --- | --- | --- | --- |
| **Dll4** | GTG CTG GGA CTG TAG CCA CT | TGT TAG GGA TGT CGC TCT CC | Flox=455bp  WT=419bp |
| **Jag1** | GCA AGT CTG TCT GCT TTA TC | AGG TTG GCC ACC TCT AAA TC | Flox=317bp  WT=267bp |
| **Cre** | TCG ATG CAA CGA GTG ATG AG | AGT GCG TTC GAA CGC TAG AG | Cre=373bp |

**Supplementary Table 2	Antibodies**

| **Antibody** | **Company** | **Use** | **Final**  **Concentration** |
| --- | --- | --- | --- |
| **Dll4** | Genentech | Blocking antibody | 10μg/ml |
| **Dll4** | Abcam (Ab7280) | WB  IF | 0.2μg/ml  1μg/ml |
| **Jag1** | R&D systems | Blocking  antibody | 10μg/ml |
| **Jag1** | Santa Cruz (sc 6011) | WB  IF | 0.2μg/ml  4μg/ml |
| **Notch1** | Abcam (ab 8925) | WB | 0.2μg/ml |
| **Notch4** | Santa Cruz (sc 5594) | WB  IF | 1μg/ml  2μg/ml |
| **CDH5** | BD Bioscience (555661) | IF | 1μg/ml |
| **pCNA** | Abcam (ab 15497) | IF | 5μg/ml |
| **Ki67** | Abcam (ab 15580) | IF | 2μg/ml |
| **CD31** | Biolegend 102513 | IF | 5μg/ml |

WB, Western blotting; IF, immunofluorescence

**Supplementary Table 3	PCR primers for real-time PCR.**

**Porcine primers:**

| **Gene** | **Forward** | **Reverse** |
| --- | --- | --- |
| **Notch1** | TGCCTGTGTCCACCTGGCTTCA | CTCCGTTTCGGCACAGGTGGGTA |
| **Notch2** | TCTGCTCACCAGGATTCA | CCTCGGGGCACATACAAC |
| **Notch3** | GCTCCTTGCCCCCACTCT | GAAACCCATTCCATCGCT |
| **Notch4** | TCCAAGAAATGCCCATAAAC | CACATAGTAGGTGCCCAATAAA |
| **Jag1** | TGTTAGCAAACGTGACGGGA | GGGGCACCAGGAAATCTGTT |
| **Dll4** | ATGCAAGAAGCGCAATGACC | CAGACAGGCTGTTCGCAGTA |
| **B2M** | TTCACTCCTAACGCTGTGGA | GTGGTCTCGATCCCACTTAAC |

**Mouse primers:**

| **Gene** | **Forward** | **Reverse** |
| --- | --- | --- |
| **Jag1** | GAGGCGTCCTCTGAAAAACA | ACCCAAGCCACTGTTAAGACA |
| **Dll4** | CGGGAACCTTCTCACTCAAC | TTGGATGATGATTTGGCTGA |
| **TBP** | GGGGAGCTGTGATGTGAAGT | CCAGGAAATAATTCTGGCTCA |

**Human primers:**

| **Gene** | **Forward** | **Reverse** |
| --- | --- | --- |
| **Notch1** | CGGGGCTAACAAAGATATGC | CACCTTGGCGGTCTCGTA |
| **Notch2** | TGGTGGCAGAACTGATCAAC | CTGCCCAGTGAAGAGCAGAT |
| **Notch3** | AGCTTGGGAAATCAGCCTTA | TCCTTGCTATCCTGCATGTC |
| **Notch4** | CCTCTCTGCAACCTTCCACT | GCCTCCATTGTGGCAAAG |
| **Jag1** | GGCAACACCTTCAACCTCA | GCCTCCACAAGCAACGTATAG |
| **Dll4** |  |  |
| **Hes1** | GCACAGAAAGTCATCAAAGCC | TTCCAGAATGTCCGCCTT |
| **MCP-1** | GCAGAAGTGGGTTCAGGATT | TGGGTTGTGGAGTGAGTGTT |
| **KLF4** | TAGCTCGAGGCATTCCAAGC | CCCGTGTGTTTACGGTAGTG |
| **HPRT** | TTGGTCAGGCAGTATAATCC | GGGCATATCCTACAACAAC |
